# Papillary and Reticular Fibroblasts Generate Distinct Microenvironments that Differentially Impact Angiogenesis

**DOI:** 10.1101/2020.11.29.402594

**Authors:** Adèle Mauroux, Pauline Joncour, Benjamin Gillet, Sandrine Hughes, Corinne Ardidie-Robouant, Laëtitia Marchand, Athanasia Liabotis, Philippe Mailly, Catherine Monnot, Stephane Germain, Sylvie Bordes, Brigitte Closs, Florence Ruggiero, Laurent Muller

**Author notes:** These authors contributed equally to this work (Florence Ruggiero, Laurent Muller).

## Abstract

Papillary and reticular dermis show distinct extracellular matrix (ECM) and vascularization, and fibroblasts isolated from these compartments have different gene expression patterns and behaviour *in vitro.* However, due to lack of relevant models, the contribution of skin fibroblast sub-populations to vascularization remains unknown. We thus cultured human papillary and reticular fibroblasts as cell sheets. Differential transcriptomic analysis was performed by RNA sequencing to characterize their microenvironment. Bioinformatic analysis revealed that each fibroblast population expressed specific angiogenesis and matrisome gene expression signatures resulting in specific ECM that differed both in composition and structure. The impact of secreted and ECM-bound factors was then assessed using 3D angiogenesis assays. When co-cultivated with endothelial cells, the papillary and reticular microenvironments induced the formation of distinct capillary networks mimicking the characteristics of vasculature of native dermis subcompartments (vessel diameter and density, number of branch points). Whereas conditioned media of papillary fibroblasts displayed intrinsic high angiogenic potential, reticular ones only contributed to capillary formation induced by exogenous VEGF. These results show that skin fibroblast populations regulate angiogenesis via both secreted and ECM-bound factors. Our work emphasizes the importance of papillary and reticular fibroblasts, not only for modelling dermis microenvironment but also for its vascularization.

## INTRODUCTION

The dermis is divided into two layers: the superficial papillary and the deep reticular dermis, which show distinct composition and specific structure of their ECM (Brown 1972; Marcos-Garcés et al. 2014; Haydont et al. 2019a; Nyström and Bruckner-Tuderman 2019). In addition, fibroblasts isolated from papillary and reticular dermis display distinct morphology, proliferation rate, and gene expression signature (Sriram et al. 2015; Lynch and Watt 2018). Each compartment display distinct functions in skin aging (Mine et al. 2008; Haydont et al. 2018), wound healing (Driskell et al. 2013), epidermis differentiation (Mine et al. 2008), basement membrane deposition (Sorrell et al. 2004; Varkey et al. 2013) and cancer progression (Hogervorst et al. 2018).

Another striking difference is that papillary and reticular dermis show distinctive vascular networks, with the upper dermis including a dense network of capillaries, terminal and post-capillary venules and the reticular dermis being crossed by fewer vessels of larger diameter (Braverman 2000; Deegan et al. 2018; Deegan and Wang 2019). Whether the dermal fibroblast sub-population and the ECM they produce can influence dermal vascularisation remains an open question. We and others suggested that papillary fibroblasts are more pro-angiogenic compared to reticular (Sorrell et al. 2008a; Nauroy et al. 2017). However, the lack of physiologically relevant *in vivo* models has limited the understanding of the molecular and cellular mechanisms underlying these processes (Philippeos et al. 2018; Zuo et al. 2019). *In vitro* models are being extensively used but mostly consist in 2D cell monolayers or 3D hydrogels made of exogenous proteins (Sorrell et al. 2004; Mine et al. 2008; Varkey et al. 2013). In addition, most of these culture systems are not appropriate for endothelial cell co-culture.

Cell sheet cultures represent a promising approach to study papillary and reticular microenvironment because they rely on the ability of cells to produce their own ECM and are more physiological (Auger et al. 2004; Pillet et al. 2017). We and others recently described that papillary and reticular fibroblasts express specific matrisome markers (Nauroy et al. 2017; Haydont et al. 2019b). Besides, cell sheets can also be used to evaluate the impact of fibroblasts microenvironment in blood vessel morphogenesis (Laschke and Menger 2016).

We thus aimed at developing vascularised papillary and reticular cell sheets to determine the contribution of the microenvironment of each sub-population of fibroblasts to capillary formation *in vitro*. We cultivated site-matched papillary or reticular fibroblasts as cell sheets in a cell medium compatible with endothelial cell culture (Gorin et al. 2016). We confirmed these cultures maintained papillary and reticular fibroblast identities. We then characterised the microenvironment and angiogenic profiles of each cell sheet by combining transcriptomic analysis, angiogenesis assays as well as TEM and SHG. We demonstrate here that papillary and reticular fibroblasts express specific angiogenesis gene signatures and that microenvironment of each fibroblast subpopulations differentially contribute to skin vascularisation.

## RESULTS

### Transcriptome of papillary and reticular cell sheets reveals differences in ECM and angiogenic gene expression

We verified the features of each population of fibroblasts for the three donors used in this study and confirmed their identity as papillary and reticular fibroblasts (Fig. S1A, S1B).

To generate a 3D microenvironment closer to dermal ECM and compatible with endothelial cell co-culture, we cultivated each population of fibroblasts as cell sheets. Using immunofluorescence and western blot, we checked for the expression of the papillary marker collagen VII and the reticular markers, elastin and transglutaminase type II (TG2) in our culture conditions (Janson et al., 2012; Nauroy et al., 2017). Reticular fibroblasts cultivated as cell sheets secreted more elastin and its expression was enriched in ECM-enriched fraction (Fig. 1A). The ECM-enriched fraction of reticular fibroblast sheets also contained higher amounts of TG2 as compared to the matched paired papillary ones (Fig. 1A). On the contrary, collagen VII was only found in the cytosolic-enriched fraction of papillary cell sheets (Fig. 1A). Proteins expression was assessed by immunofluorescence and consistent with western blot analysis. (Fig. 1B). We concluded that cell sheet culture permitted to maintain fibroblast sub-population phenotypes in long-term culture.

**Figure 1:**
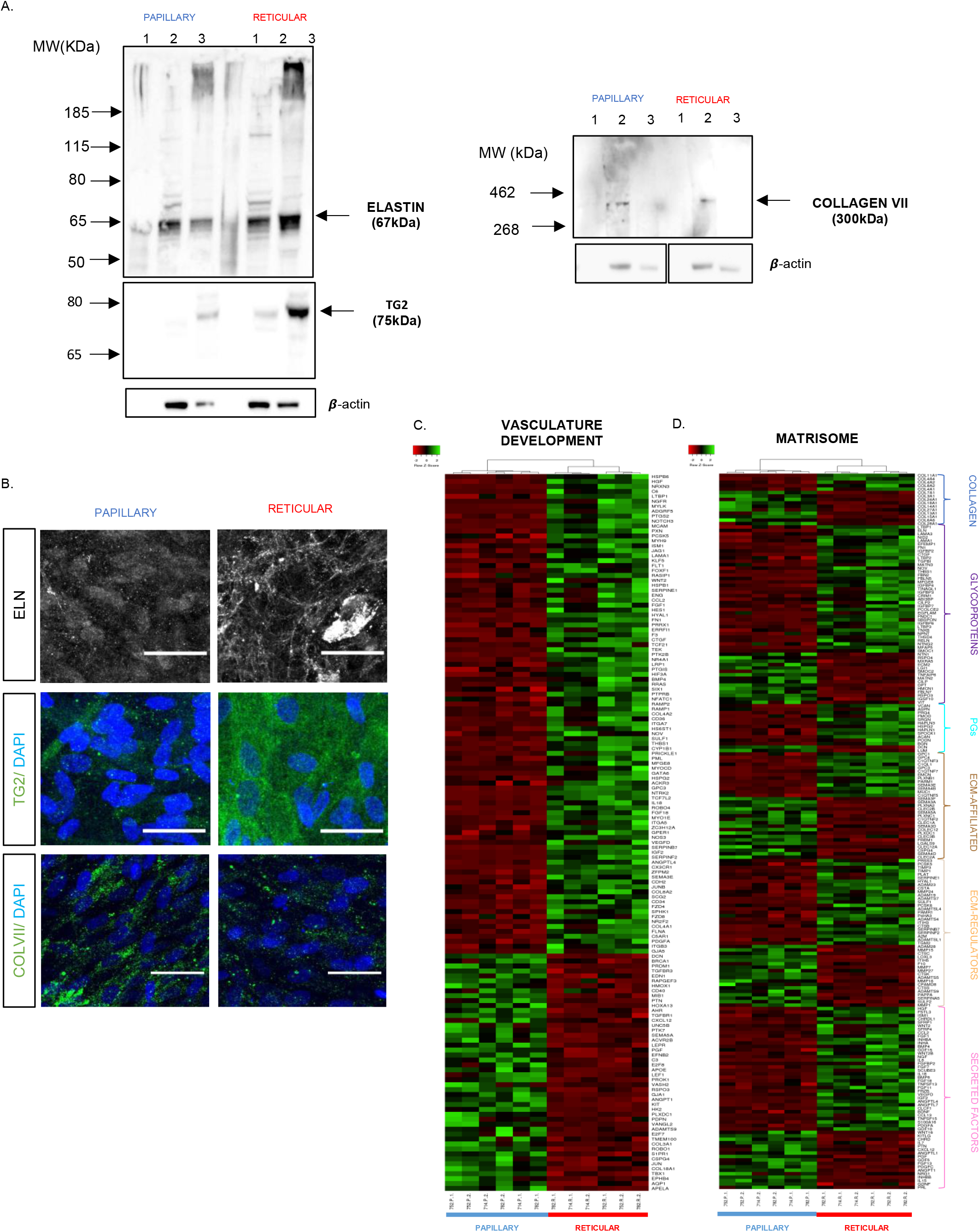
Cell sheet culture preserves papillary and reticular fibroblasts identity and show distinct matrisome and angiogenic gene signature. **A.** Matrisome markers of reticular identity (elastin, transglutaminase type II) and papillary identity (collagen VII) were immunodetected in distinct cell sheets fractions by western blot. Lane 1, 2 and 3 correspond respectively to secretion medium precipitate, cytosolic-enriched and ECM-enriched fractions. **B.** Immunofluorescences for markers of papillary and reticular matrisome were also performed on cell sheets after two weeks of culture. ELN : elastin; TG2 : type II transglutaminase ; COLVII : collagen VII. Scale bar: 200 μm. **C.** Heatmap of differentially regulated genes of the GO term “Vasculature Development” analyzed by hierarchical clustering. FC≥1.5, p<0.05. **D.** Heatmap of differentially regulated genes in papillary and reticular matrisome analyzed by hierarchical clustering. FC≥1.5, p<0.05.

We then performed RNAseq analysis of matched pairs of papillary and reticular fibroblasts after one week of cell sheet culture. At this time-point fibroblasts already organized into multilayers and actively produced and remodelled their microenvironment. We performed a Gene Ontology (GO) analysis of biological processes and found an enrichment of GO terms in papillary and reticular cell sheets that was consistent with previous studies (Janson et al. 2012; Nauroy et al. 2017; Philippeos et al. 2018) (Fig. S3C and D, Table S1).

Interestingly, the GO term related to “Sprouting angiogenesis” was enriched in papillary fibroblast sheets whereas reticular fibroblast sheets displayed a more diverse panel of GO terms related to blood vessel formation (Table S2). We further characterised the angiogenic gene expression profiles of papillary and reticular fibroblast sheets by listing all the genes belonging to the GO term “Vasculature development” (Table S3) and observed a clustering of samples according to fibroblasts subtypes, indicating specific angiogenic expression signature (Fig 1C).

GO terms related to ECM synthesis and organization were enriched in reticular fibroblast sheets, while those related to ECM degradation were enriched in papillary ones (Table S2). We characterised the matrisomes of both fibroblast sheets as described by Naba et al. (Naba et al. 2012) (Table S4). As for angiogenic gene expression profiles, papillary and reticular fibroblast sheets showed specific matrisome gene expression signatures (Fig. 1D).

### Paracrine regulation of angiogenesis by papillary and reticular cell sheets

To further characterize the angiogenic gene expression signatures of papillary and reticular fibroblast sheets, we extracted genes of the GO term “Vasculature development” that encoded secreted diffusible factors (Table S5, FC ≥ 2, Mean TPM ≥ 10). Interestingly, reticular fibroblasts expressed a balance of pro-, anti- and context-dependent regulators of angiogenesis, while papillary fibroblasts only expressed pro-angiogenic factors (Fig. 2A).

**Figure 2:**
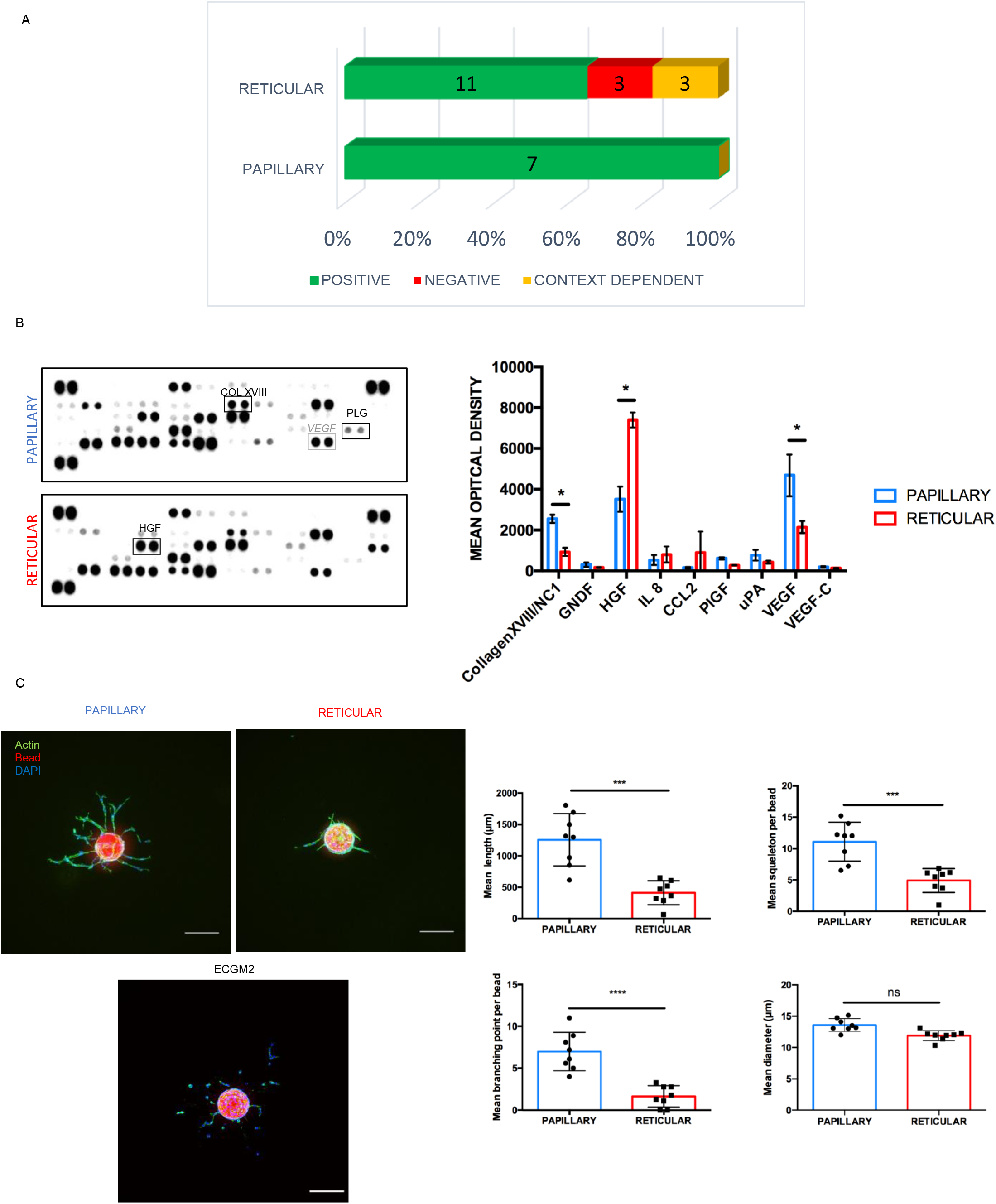
Papillary fibroblasts have a higher angiogenic potential compared to reticular. **A.** Distribution of genes coding for secreted pro- anti- or context dependent regulators of angiogenesis. **B.** Analysis of papillary and reticular conditioned media by antibody array. Proteins identified in transcriptomic study are indicated by black frames. Gray frames indicate other identified secreted proteins. Right panel : Quantification of mean optical density for all three donors (FC≥1.5). Mean ± SD, T-Test, * : p<0.05. **C.** Representative images of capillary formation in a 3D fibrin hydrogel assay using papillary or reticular conditioned media. Scale bar: 200 μm. The length, vessel number, branching points and diameter were quantified for each condition. Graphs represents the mean of two independent experiments carried out using three donors and different batches of conditioned media (n=8). Mean ± SD; T-test, *** p=0.001, **** : p<0.001.

Using an antibody array, we next verified the expression of selected secreted factors at the protein level. (Fig. 2B). High levels of HGF were found in reticular fibroblast sheet conditioned media, confirming differences of gene expression observed in our transcriptomic study. Likely, high levels of XVIII was detected in the papillary conditioned media as Placental Growth Factor (PlGF) even if this difference was not significant. In addition, papillary fibroblasts secreted higher level of VEGF compared to the reticular one.

We then assessed the effect of these conditioned media on vessel formation using 3D fibrin gel-based angiogenic assay. Papillary conditioned media promoted the formation of three times longer capillary sprouts than reticular one. This was due to an increase in both the number of primary capillary sprouts from the beads and the number of branching points by 2.26 and 5.83 folds, respectively (Fig. 2C). Addition of the pro-angiogenic factor VEGFA clearly stimulated capillary formation induced by both fibroblasts conditioned media (Fig. S4). Even if factors secreted by reticular fibroblasts were less efficient in stimulating sprouting angiogenesis than site-matched papillary fibroblasts, they did promote the formation of capillaries compared to endothelial cell media only (Fig. 2C).

Overall, these results indicated that papillary fibroblast sheets have a higher angiogenic potential compared to site-matched reticular cell sheets due to distinct angiogenic gene expression profiles and secretome.

### Papillary and reticular cell sheet microenvironment exhibit distinct composition and ultrastructure

We further characterised the matrisome of papillary and reticular cell sheets by extracting the matrisome data from each cell sheets and creating subsets of genes coding for proteins involved in distinct biological processes based on the literature (Fig. 3A). Distribution of matrisome genes between these defined categories revealed that immunity, basement membrane zone components and proteases gene sets were enriched in papillary cell sheet matrisome whereas genes upregulated in reticular matrisomes encoded regulators of elastic fiber assembly and fibrillary collagen biosynthesis and fibrillogenesis (Fig. 3A).

**Figure 3:**
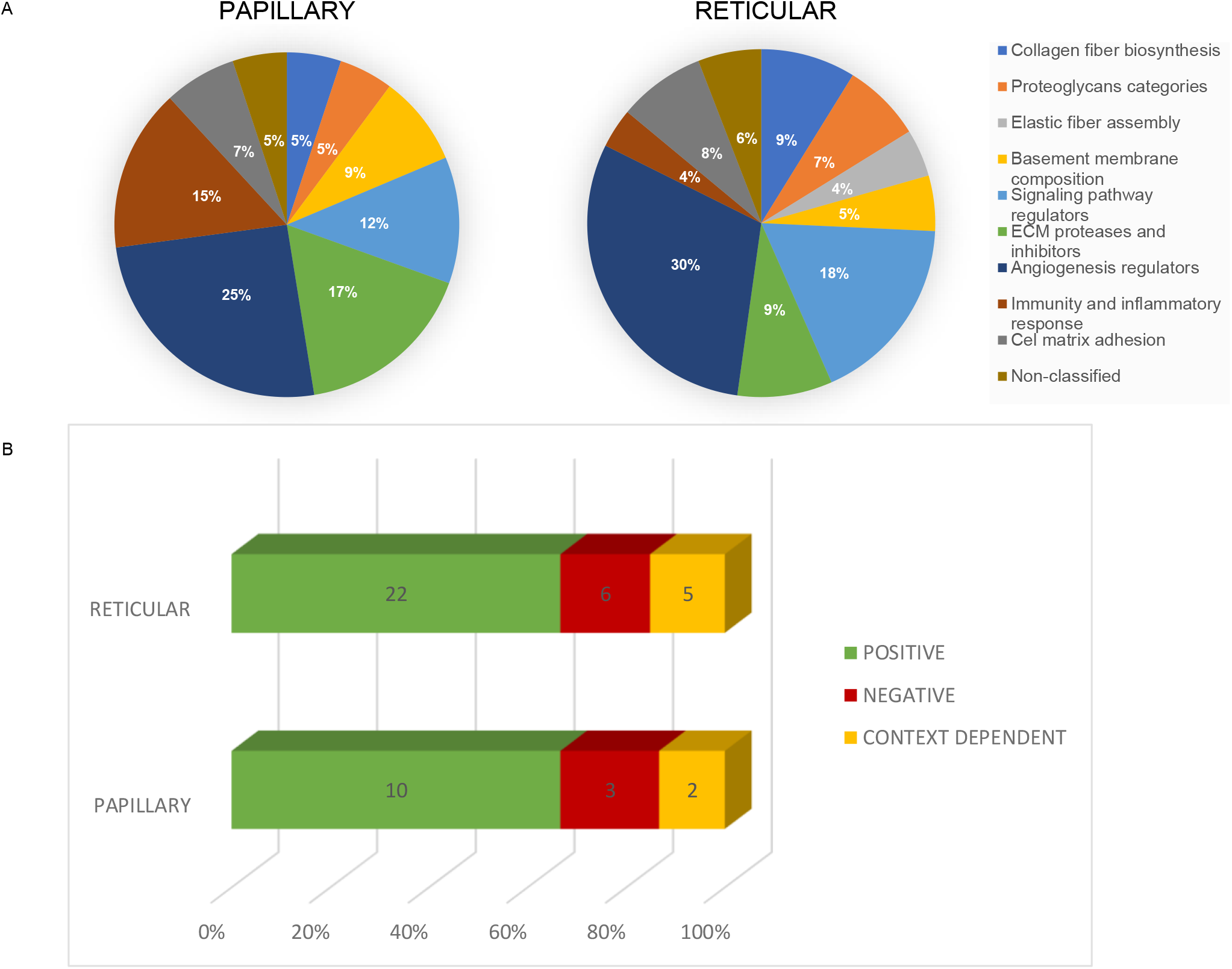
Papillary and reticular cell sheets matrisome suggest the organization of distinct microenvironment. **A.** Distribution of matrisome genes between functional categories based on literature. The percentage of genes represented for each sub-categories are indicated on the diagram. **B.** Distribution of matrisome genes coding for regulators of angiogenesis classified as pro- anti- or context-dependent.

Only matrisome of reticular cell sheets comprised genes involved in elastic fiber assembly (*ELN, FBN2, MAFP5, TNXB*) and expressed a large diversity of proteoglycans (Table S7) (Table 3). Reticular sheets also expressed genes regulating fibrillar collagen biosynthesis and fibrillogenesis (*CTGF, TGFBI, BGN, TNXB*). The matrisome of papillary cell sheet comprised genes encoding unconventional collagens present at the dermo-epidermal junction (*COL7A1, COL6A6, COL18A1, COL15A1*). We found that papillary and reticular cell sheets both expressed basement membrane tool kit (Hynes 2012). Notably, reticular fibroblasts highly expressed *COL4A1* and *COL4A2*. Papillary cell sheets expressed a higher ratio of proteases over protease inhibitors compared to the reticular ones, suggesting a higher ability to remodel dermal ECM (Table S6).

Interestingly, papillary and reticular cell sheets expressed matrisome genes coding for positive, negative and context-dependent regulators of angiogenesis in similar proportion as compared to secreted factors only (Fig. 3B, Table S8). Pro-angiogenic factors of the papillary matrisome mainly consisted of guidance molecules (*SEMA5A, SEMA4D, NTN1, PLXDC1*) while reticular cell sheets mostly expressed ECM structural component genes (*COL4A1, COL4A2, COL8A2, LAMA1, EFEMP1*). Negative regulators of angiogenesis included ECM remodeling enzymes, as *TGM2* and *SULF1*, whereas negative regulators present in papillary matrisome were genes encoding for semaphorin guidance molecules. Finally, context-dependent regulators of angiogenesis of the papillary matrisome were the multiplexins *(COL15A1, COL18A1)* and those in reticular cell sheets were gene coding for member of the angiopoietin-like family *(ANGPTL4, ANGPTL7)*.

We next characterised the structure and organisation of cell sheet ECM using Second Harmonic Generation two-photon microscopy. Interestingly, collagen fibres in reticular ECM were overall more aligned compared to papillary ECM as indicated by the distribution of collagen fibres directionality (Fig. 4A). These results are representative of an experience gathering three matched paired donors. We further characterised the ECM by TEM. Strikingly, reticular cell sheet ECM consisted of aligned collagen fibres that often formed large bundles whereas papillary cell sheet ECM contained loosely organized collagen fibrils (Fig. 4B vii-x), confirming SHG observations. Deposition of electron dense patches of elastin was observed in the reticular cell sheets (Fig. 4B viii). In papillary cell sheets, ECM was mainly deposited as thin unstriated fibrillar material (Fig. 4B, v) and contained electron-dense proteoglycan aggregates (Fig. 4B, vii). The ECM structure and organization of the fibroblast subpopulations were consistent with their matrisome signatures.

**Figure 4:**
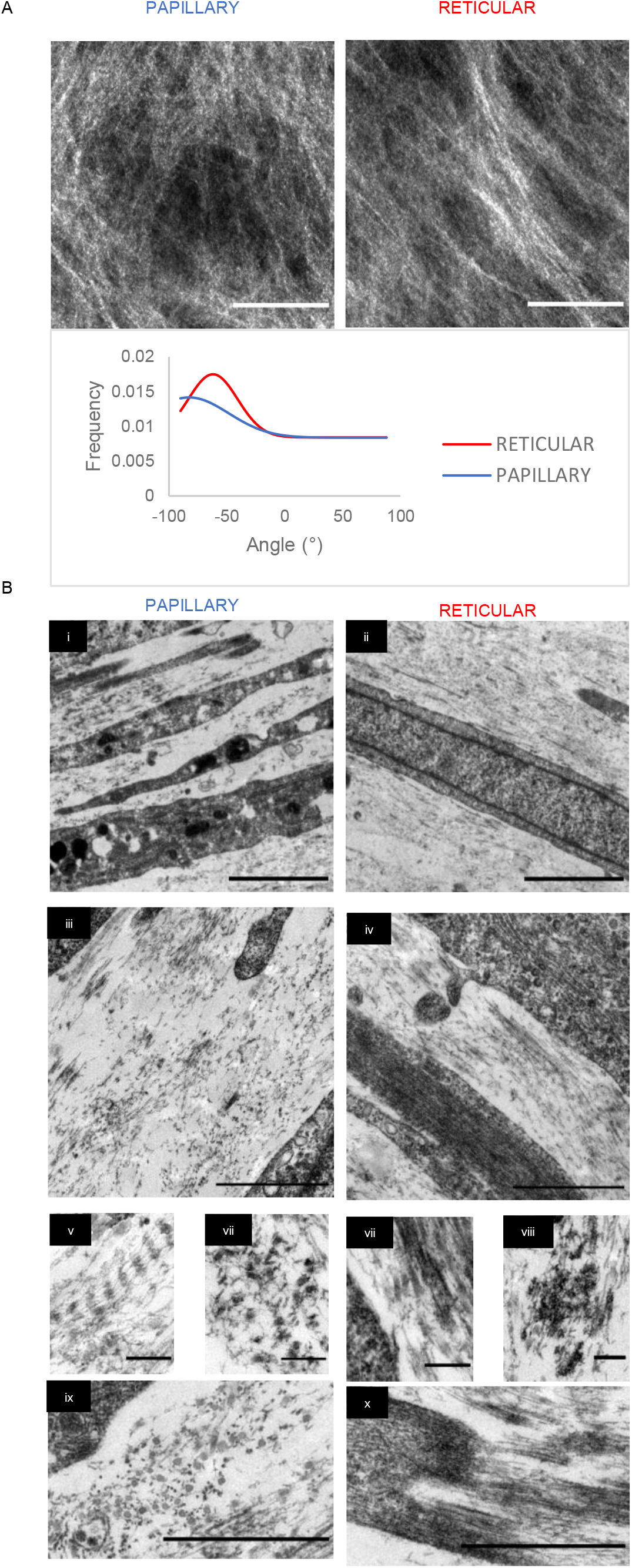
Papillary and reticular cell sheets have distinct ECM structure and organization. **A.** Representative images of papillary and reticular collagen fiber organization using SHG. Summed projection, Scale bar: 50 μm. Representative fiber directionality plot showing the distribution of fibers as function of angles. N=1 **B.** Papillary and reticular cell sheets ECM ultrastructure after three weeks of culture observed by TEM. Representative images show a greater alignment of fibers in the reticular cell sheets (**ii**; **iv**) compared to papillary (**i**; **iii**) Scale bars: 2 (**I** and **iii**) and 1 (**ii** and **iv**) μm. (**v**-**x**) Magnification of cell sheets ECM. Immature collagen fibrils (**v**), interstitial non-fibrillar material (**vii**) and collagen fibers in papillary ECM (**ix**). Representative aligned collagen fibers (**vii**, **x**) and electronic dense elastin aggregates (**viii**) in reticular ECM. Scale bar: 200 nm.

### Papillary and reticular cell sheet microenvironments differentially influence capillary formation

We then assessed the effect of papillary and reticular cell sheet microenvironments on capillary formation using a co-culture angiogenesis assay. In this model, capillaries were embedded within the fibroblast cell sheet ECM (Fig. 5A). Endothelial cells organised into mature capillaries stabilised by a basement membrane (Fig. 5B). We co-cultivated HUVECs onto papillary or reticular fibroblasts cell sheets for a week in the presence or absence of VEGF and characterised the vascular network formed within both cultures using the endothelial marker CD31 (Fig. 5C). Without VEGF, both papillary and reticular microenvironments were able to elicit capillary formation. Interestingly, the mean diameter of capillaries formed in the reticular microenvironment was significantly larger compared to the papillary one (Fig. 5C, zoom and statistics). However, in the presence of VEGF, the papillary cell sheet supported the formation of a denser and longer vascular network as verified by quantification (Fig. 5C). The vascularised area was 20% larger in papillary microenvironment compared to the reticular one and resulted in a longer and more branched network. Papillary microenvironment supported the formation of a plexus-like denser network of micro-vessels whereas reticular microenvironment induced the formation of a few aligned vessels of larger diameters.

**Figure 5:**
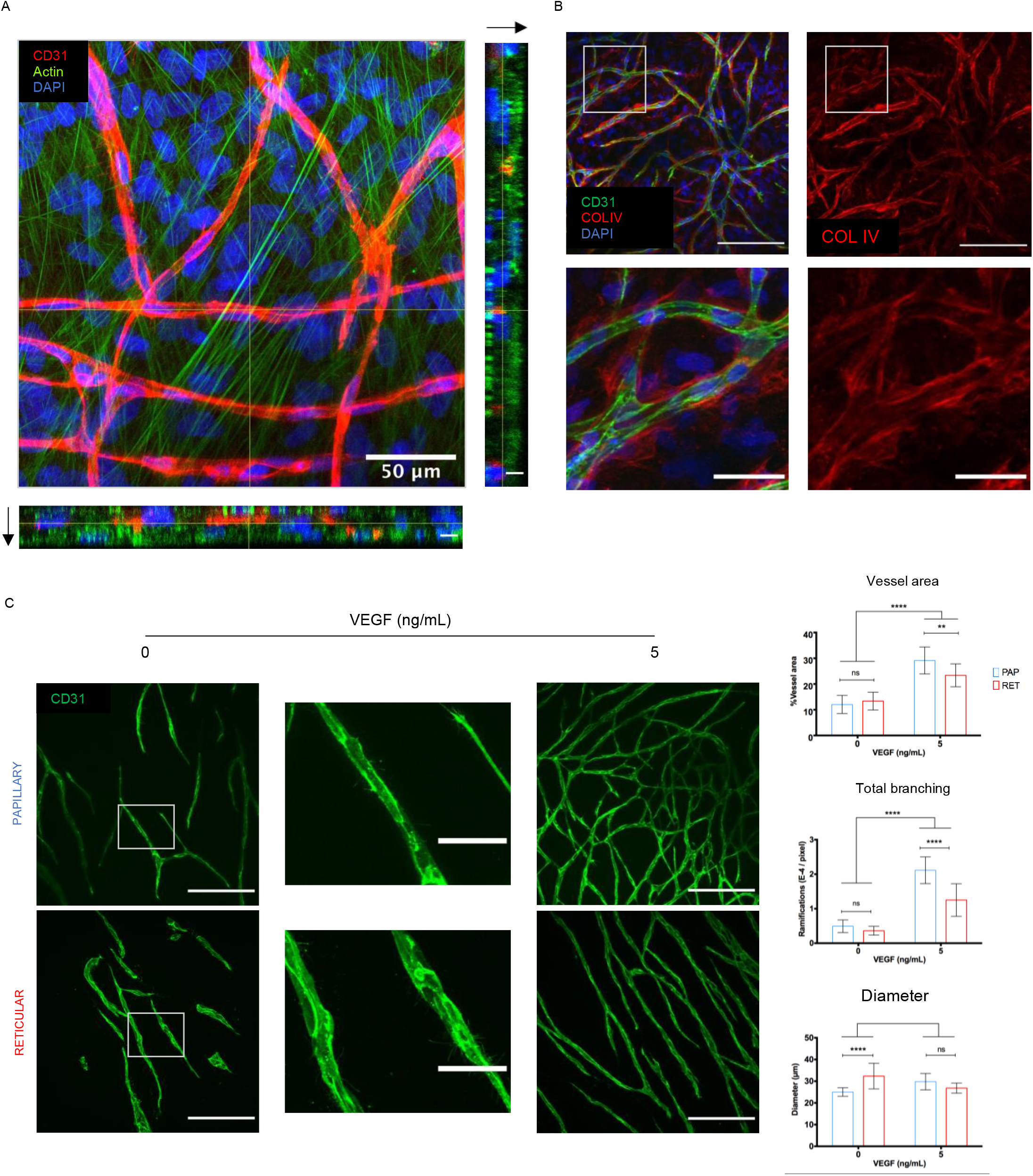
Papillary and reticular cell sheets microenvironment differentially affect capillary formation. **A.** Staining of cytoskeleton and vascular marker CD31 indicates that capillaries are embedded within the microenvironment generated by fibroblasts. Maximum intensity projection and corresponding orthogonal views. Scale bar: projection 50 μm; orthogonal views 10 μm. **B.** Capillaries are stabilized by a basement membrane detected by collagen IV immunofluorescence. Maximum Intensity Projection. Scale bar: 200 μm; zoomed area: 50 μm. **C.** Representative images of vascular network formed in papillary or reticular cell sheet microenvironment. CD31 staining. Scale bar: 200 μm; zoomed area: 50 μm. Corresponding quantification are reported on the right hand side. Graph represents the mean of two independent experiments carried out using three donors in triplicates (n=6). Mean ± SD; 2 way ANOVA; **: p<0.01; ****: p<0.001.

## DISCUSSION

In this study, we showed that skin fibroblast populations specifically contribute to angiogenesis regulation. Cell sheet culture reproduced some features of native ECM dermal compartments and recapitulated the main characteristics of skin vascularisation: dense and branched networks of capillaries in the papillary dermis, and a few vessels with larger diameter in the reticular dermis. These differences resulted from distinct paracrine and ECM-dependent mechanisms due to specific gene expression signatures.

The higher angiogenic potential of papillary fibroblasts was associated with a pro-angiogenic gene expression signature. Papillary conditioned medium was enriched in VEGF and PlGF that are positive regulators of skin angiogenesis (Romer et al. 1996; Detmar 2000; Dewerchin and Carmeliet 2012; Rubina et al. 2017). Even though we detected high levels of secreted HGF, the angiogenic potential of reticular fibroblasts conditioned medium was lower compared to papillary. The co-expression of genes coding for anti-angiogenic proteins might have limited angiogenesis induced by reticular fibroblast conditioned media. VEGFA was previously found to be secreted in both papillary and reticular fibroblasts at equal levels, quantification of VEGFA by ELISA would allow to appreciate the difference with our study (Mine et al. 2008; Sorrell et al. 2008b). Increased secretion of CCL2/MCP1 by reticular fibroblasts was previously described (Mine et al. 2008), whereas HGF was reported as upregulated in papillary fibroblast cultures (Sorrell et al. 2008b). Cell sheet culture was likely responsible for these differences.

Contrary to paracrine assay, capillary formation in papillary and reticular microenvironment was identical in the absence of VEGF. This may results from the sequestration of growth factors by the ECM and/or non-soluble ECM components. This hypothesis is supported by the equal distribution of pro-, anti- and context-dependent matrisome regulators of angiogenesis between papillary and reticular cell sheets. Reticular microenvironment induced the formation of larger capillaries than papillary fibroblasts in absence of exogenous VEGF. Addition of VEGF in our culture system is likely to have erased this subtle difference (Nakatsu et al. 2003).

Papillary microenvironment was more responsive to VEGFA-165 compared to reticular microenvironment. This is of particular interest as superficial vascular network is the most remodelled during wound healing and is altered in chronic inflammatory disease. Papillary fibroblast sheets promoted the formation of a plexus-like vascular network that is consistent with their semaphoring gene expression profile (Adams and Eichmann 2010). As previously described (Janson et al., 2012), TG2 was found enriched in reticular fibroblasts cell sheet ECM. With *SULF1*, these remodelling enzymes might contribute to the reduced capillary formation in reticular microenvironment (Narita et al. 2006; Beckouche et al. 2015).

We also demonstrated that papillary and reticular fibroblast sheets produce an ECM that displayed similar organisation than in native skin. Reticular microenvironment contained mature elastic fibers. Reticular fibrillar network was also more structured than the papillary one as previously observed in skin biopsies and decellularised ECM (Marcos-Garcés et al. 2014; Ghetti et al. 2018). ECM topography influences endothelial cell adhesion and capillary formation, branching and diameter (Du et al. 2014; McCoy et al. 2018). These differences in ECM organisation and structure might thus also contribute to the distinct angiogenic properties of each fibroblasts populations.

Recent studies have identified cell-surface markers of papillary and reticular fibroblasts that were used to isolate the sub-population of fibroblasts using FACS (Philippeos et al. 2018; Korosec et al. 2019). It would thus be interesting to generate cell sheets using cell-sorted papillary and reticular fibroblasts based on the expression of these markers and to assess if they display similar microenvironment and angiogenic properties identified in the present study. It will also be important to assess the angiogenic properties of papillary and reticular fibroblasts on site-matched endothelial cells human dermal microvascular endothelial cells (HDMEC).

Vascularised papillary and reticular cell sheets represent an appealing model to study the impact of fibroblasts microenvironment in skin biology such as vascular aging (Li et al. 2006; Gunin et al. 2011) or lymphangiogenesis (Skobe and Detmar 2000). They could also be used to generate vascularised micro-tissues by cell sheet layering (Gibot et al. 2010; Cerqueira et al. 2014). Lastly, the therapeutic potential of vascularised papillary and reticular cell sheets could be assessed in regenerative medicine to treat chronic wounds.

Overall, our study proposed angiogenic gene expression signature profiles for the two fibroblast sub-populations including secreted angiogenic factors and matrisome genes. These results emphasise the importance of the microenvironment composition and structure in angiogenesis regulation. Moreover, vascularised papillary and reticular cell sheets developed and characterized in this study could be used in the next future for bioactive molecule screening and regenerative medicine applications.

## MATERIAL AND METHODS

For more details, see supplemental Methods.

### Cell culture

Human umbilical venous endothelial cells (HUVEC) were prepared and cultured as previously described (Chomel et al. 2009). HUVEC between passage 2 and 4 were used. Papillary and reticular fibroblasts were isolated and amplified as previously described (Nauroy et al. 2017). Briefly, site-matched papillary and reticular dermis were separated mechanically using a dermatome and explants from each compartments put in culture and amplified. All experiments were carried out using three healthy female donors aged from 30 to 50 who went through mammal or abdominal surgery and gave their written consent. Site-matched papillary and reticular fibroblasts were used between passages 5 to 8.

### Papillary and reticular cell sheet culture

Site-matched papillary and reticular fibroblasts were seeded at 20 000 cells/cm^2^ in endothelial cell culture medium ECGM2 (Promocell, Heilderberg, Germany) depleted in basic Fibroblasts Growth Factor (bFGF) and Vascular Endothelial Growth Factor A (VEGFA) (ECGM2ΔFGFΔVEGF) supplemented with Pennicilin/Streptomycin (Life Technologies, Carlsbad, CA, USA). Cell medium was changed three times a week. After one week, the culture consisted of a multilayer of cells embedded in an ECM that we refer as cell sheet. For matrisome papillary and reticular markers cell sheets were cultivated for two weeks, for ANGPTL4 immunofluorescence cells were cultivated from one to three weeks.

### Transcriptomic analysis using RNA sequencing

RNA sequencing experiments were performed by the IGFL sequencing platform (Lyon, France). Site-matched papillary and reticular fibroblasts of the three donors used in the study were cultured as cell sheets for one week. Briefly, RNA was extracted and concentrations and quality of RNA assessed. Cell sheets transcriptomes were sequenced by Illumina NextSeq^®^500 Wide Genome Sequencing system (Illumina, San Diego, CA, USA). Statistical and bioinformatical analyses were performed using R software and DEseq2 statistical analysis (R Core team (2013)). Genes were reported as differentially expressed when fold change was superior to 1.5 and p-value≤0.05. Gene Ontology (GO) term analysis was carried out using DAVID Bioinformatics Ressources 6.8 (Huang et al. 2009a, 2009b). Angiogenic and matrisome genes were obtained by extracting genes belonging to the GO term “Vasculature Development” (GO:0001944) and to the human matrisome database (Naba et al. 2016). Heatmap and hierarchical clustering analyses were done using Heatmaper (Babicki et al. 2016).

### Western Blot

Papillary or reticular fibroblasts were cultivated as cell sheets for one week. Proteins secreted overnight were precipitated with Trichloroacetic acid (TCA) (10%). Cell sheet lysates were collected using NP40/DOC lysis buffer as previously described (Bignon et al. 2011) and both supernatant (cytosolic fraction) and pellets (ECM-enriched fraction) were analyzed. Proteins were separated by SDS-PAGE and transferred on a PVDF membrane from pre-casted gels (Invitrogen, Carlsbad, CA, USA). Primary antibodies were diluted in 5% non-fat milk. Immunological detection was performed using horseradish peroxidase secondary antibodies and ECL or Femto ECL kit (Invitrogen, Carlsbad, CA, USA). Images were acquired using Fusion Capt Advance FX7 (Vilber, Eberhardzell, Germany).

### Conditioned medium (CM) preparation

Papillary or reticular fibroblasts were seeded at 20 000 cells/cm^2^ in 75 cm^2^ flasks in basal medium. Cells were grown until reaching confluence and medium was switched to ECGM2 and renewed every day for seven days. CM was harvested every day and filtered through 0.22 μm filter. CM from the first 24 hours were discarded and CM from day four to seven pooled, aliquoted and stored at −80°C.

### Antibody array

Conditioned media of papillary and reticular fibroblasts from the three donors were analysed using human angiogenesis antibody array (R&D systems, Minneapolis MN, USA). Images were acquired using Fusion Capt Advance FX7 (Vilber, Eberhardzell, Germany).

### 3D capillary formation assay

3D angiogenesis assay in fibrin gel was carried out and analysed as previously described (Bignon et al. 2011; Gorin et al. 2016; Gaborit et al. 2019). Briefly, Cytodex^®^ microcarrier beads (SIGMA-Aldrich, Saint-Louis, MI, USA) were covered with HUVECs and embedded 24 hours later in a 2,5 mg/mL fibrin gel in the presence or absence of VEGFA-165 at 5 ng/mL (R&D systems, Minneapolis MN, USA). Conditioned media from papillary or reticular fibroblasts were added on top of the gel and renewed every day for four days. Cultures were fixed by 4% PFA and the cytoskeleton was labelled using the antibody phalloidïn conjugated with Alexa Fluor 488 (Thermofisher, Whaltham, MA, USA). Nuclei were stained using DAPI 1/20 000. Images were acquired with Zeiss Axiozoom Apotome 2 (Zeiss, Oberkochen, Allemagne) using an automatic acquisition plugin. Quantification was performed using a plugin previously developed in the laboratory to quantify the number of capillary sprouts, total branch length, branching points and mean capillary diameter per bead.

### Fibrillary collagens imaging using Second Harmonic Generation

Cell sheets were cultivated for three weeks to maximise ECM deposition and maturation. SHG signal was acquired using upright SP5 scanning confocal microscope (Leica biosystems, Nussloch, Germany) coupled with high pulse multiphoton laser Mai Tai^®^ Ti:Sapphire (MKS Spectra Physics, Andover, Ma, USA). Samples were excited at 880 nm and SHG signal was collected at 440 nm. Z-stacks of all cell preparations were acquired and z projection of SHG signal were analysed using the directionality plugin in Fiji (Liu 1991).

### Transmission electron microscopy

To study the ECM ultrastructure, papillary or reticular cell sheets were cultivated for 3 weeks. Cultures were fixed by 2% PFA and 2.5% glutaraldehyde diluted in 0,2M sodium cacodylate at pH 7.42 for two hours at room temperature. Four mm punches of cell sheets were harvested and post-fixed in 1% osmium tetraoxide. Cell sheets were gradually dehydrated in successive baths of ethanol and included in Epon resin. Ultrathin slices of papillary and reticular cell sheets were contrasted in 7% uranyl acetate and lead citrate and imaged using Philips CM120 transmission electronic microscope.

### Co-culture angiogenesis assay

After one week of culture, papillary or reticular cell sheets were seeded with HUVECs at 6 000 cells /cm^2^ in ECGM2ΔFGFΔVEGF supplemented or not with VEGF (R&D systems, Minneapolis MN, USA) at 5 ng/mL. After one week, cell cultures were fixed with 4% PFA and vascular network detected by CD31 (see immunofluorescence) To quantify the vascular network, 50% well surface was imaged using Zeiss Axiozoom Apotome 2 (Zeiss, Oberkochen, Germany). Vascular network was quantified using a plugin developed in the laboratory to calculate total branch number, vascularised area, diameter and vascular length. Briefly, after automatic mask generation, the vascular network was skeletonized and quantified using Analyze skeleton library.

### Immunofluorescence

Cell sheets and co-culture were fixed by 4% PFA and permeabilised with Triton X-100 (0.5%). Primary and secondary antibodies used in this study are listed in supplementary data. Nuclei were labelled using DAPI 1/20 000. Images were acquired using Spinning disk W1 confocal microscope and Zeiss Observer Apotome 2 (Zeiss, Oberkochen, Germany).

### Statistics

After verifying that the data followed a normal distribution, statistical analyses were performed using i) unpaired T-Test for the 3D angiogenesis assay in fibrin hydrogel, ii) Two-way ANOVA for the quantification of the vascular network in papillary and reticular cell sheet microenvironments. (GraphPad Prism 6, GraphPad Software, La Jolla, CA, USA).

## CONFLICT OF INTEREST

AM, LM, SB, and BC are employees of SILAB. FR and LM declare the receipt of a grant from SILAB. PJ, BG, SH, CA, AL, PM, CM and SG state no conflict of interest

## SUPPLEMENTARY MATERIAL

### Proliferation

Experimental duplicates were included for each population of fibroblasts. Site-matched papillary and reticular fibroblasts were seeded at 3 300 cells/cm^2^ in 175 cm^2^ flasks in MEM supplemented with 10% SVF (Life Technologies, Carlsbad, CA, USA) and 50 U/mL of Pennicilin/Streptomycin (Life Technologies, Carlsbad, CA, USA). Fibroblasts were cultivated until the first population of fibroblasts reached 90% confluence for at least 3 passages. Population doubling time was then calculated and doubling times were normalised to papillary fibroblasts values.

### Papillary and reticular cell culture morphology

Site-matched papillary and reticular fibroblasts were cultivated as isolated cells (3 000 cells/cm^2^) in fibroblasts basal medium (MEM-10%SVF) on Ibidi 8 well slides (Ibidi, Gräfelfing, Allemagne) coated or not with rat tail collagen type I (33.5 μg/mL in 20 mM acetic acid) (BD biosciences, Franklin Lakes, NJ, USA) or plasmatic fibronectin (kindly provided by Dr. Pauthe, at 10 μg/mL). After 24 hours, cultures were fixed by 4% PFA. The cytoskeleton was labelled using phalloidin conjugated with Alexa Fluor 488 (Thermofisher, Whaltham, MA, USA) and focal adhesions were detected using anti-vinculin antibody (SIGMA V9131, 1/400). Secondary anti-mouse antibody coupled with Alexa Fluor 555 (Thermofisher, Whaltham, MA, USA) was used to detect anti-vinculin antibody and diluted at 1/1000 and nuclei were labelled with DAPI 1/20 000. Images were acquired using Zeiss Observer Apotome 2 (Zeiss, Oberkochen, Allemagne).

### Transcriptomic analysis

RNA was extracted using Nucleospin RNA kit (Macherey-Nagel, Düren, Germany) according to manufacturer instructions. RNA concentrations were measured by Qubit^®^ 2.0 fluorometer (Thermofisher, Whaltham, MA, USA). RNA quality was verified on Tapestation 2 200 using RNA screentape assay (Agilent technologies company, Santa Clara, CA, USA). All samples had a RNA Integrity Number (RIN) superior to 9.6. mRNA barcode libraries were built using SENSE mRNA-Seq Library Prep kit V2 (Lexogen, Greenland, NH, USA). Cell sheets transcriptomes were sequenced by Illumina NextSeq^®^500 Wide Genome Sequencing system with a 75bp single end run (Illumina, San Diego, CA, USA). Reads were then mapped on the human genome with RSEM (Li et al., 2011) to obtain raw read counts and Transcripts Per kilobase Million (TPM). Statistical and bioinformatical analyses were performed using R software (R Core team (2013)). Principal Composent Analysis (PCA) was performed on TPM for all sequenced genes with FactoMineR package (Lê et al., 2008). Only protein coding genes were analysed in this study. Based on the distribution of TPM in all samples, only genes that had a TPM≥20 for at least two samples were considered as expressed. Differential expression analysis was carried out using DESeq2 package (Love et al., 2014). Genes were reported as differentially expressed when fold change was superior to 1.5 and p-value≤0.05. Gene Ontology (GO) term analysis was carried out using DAVID Bioinformatics Ressources 6.8 (Huang et al. 2009a, 2009b). Only GO terms that had a Fold Enrichement (FE) of at least three and contained a minimum of 5 genes were studied. To study the angiogenic and matrisome profile, the list of all differentially regulated genes was compared to the genes belonging to the GO term “Vasculature Development” (GO:0001944) and to the human matrisome database (Naba et al. 2016). Heatmap and hierarchical clustering analyses were done using Heatmaper (Babicki et al. 2016). To identify specific gene signatures, a higher fold change was used, FC≥2, with a minimum mean TPM of 10. Genes coding for secreted factors were extracted from the list of genes belonging to the GO term “Vasculature development”. Genes were classified as pro-, anti- or context-dependent regulators of angiogenesis based on the literature. Sets of genes were created from the matrisome of papillary and reticular cell sheets based on our transcriptomic data. For the matrisome angiogenesis category, matrisome genes belonging to the GO term “Vaculature Development” were extracted. In all tables, the level of expression is reported as low (TPM≤60), medium (60<TPM<1000) or high (TPM≥1000).

### Immunofluorescence

Table of primary antibodies used in this study

| Antibody | Supplier | Species | Dilution |
| --- | --- | --- | --- |
| Anti-CD31 | M0823 DAKO | MOUSE | 1/100 |
| Anti-COLIV | 20411 NOVOTEC | RABBIT | 1/250 |
| Anti-collagen VII | C6805 SIGMA | MOUSE | 1/100 |
| Anti-elastin | 25013 NOVOTEC | GUINEA PIG | 1/200 |
| Anti-TGM2 | Ab2386 | MICE | 1/400 |
| Anti-ANGPTL4 Cterm | 6G11 HOME-MADE | MICE | 1/100 |
| Anti-collagen III | Ab7778 ABCAM | RABBIT | 1/500 |

Secondary anti- mouse, - rabbit and -guinea pig antibodies conjugated with Alexa Fluor 488, 555 or 647 were used (Invitrogen, Carlsbad, CA, USA).

### Western blot analysis (complement)

Pre-casted gels were Bolt™ 4-12% Bis-Tris gels (Invitrogen, Carlsbad, CA, USA) or NuPAGE™ 3-8% Tris-Acetate gel (Invitrogen, Carlsbad, CA, USA). Primary antibodies used in the western blot analysis were anti-collagenVII/NC1 antibody (kindly provided by Dr. Koch E., 1/500), anti-elastin (NOVOTEC 25 011, 1/2000), anti-TG2 (ABCAM ab2386, 1/2000) and anti-ANGPTL4 C-terminal fragment (6G11 antibody, 1/100).

### Co-culture angiogenesis (complement)

To evaluate capillary position within the culture, cell cytoskeletons were detected using phalloïdin conjugated with Alexa –Fluor 488 (Thermofisher, Whaltham, MA, USA; 1/400) and images were acquired using Leica SP5 inverted scanning confocal microscope (Leica biosystems, Nussloch, Germany). Orthogonal projections were generated with Fiji.

**Figure S1 :**
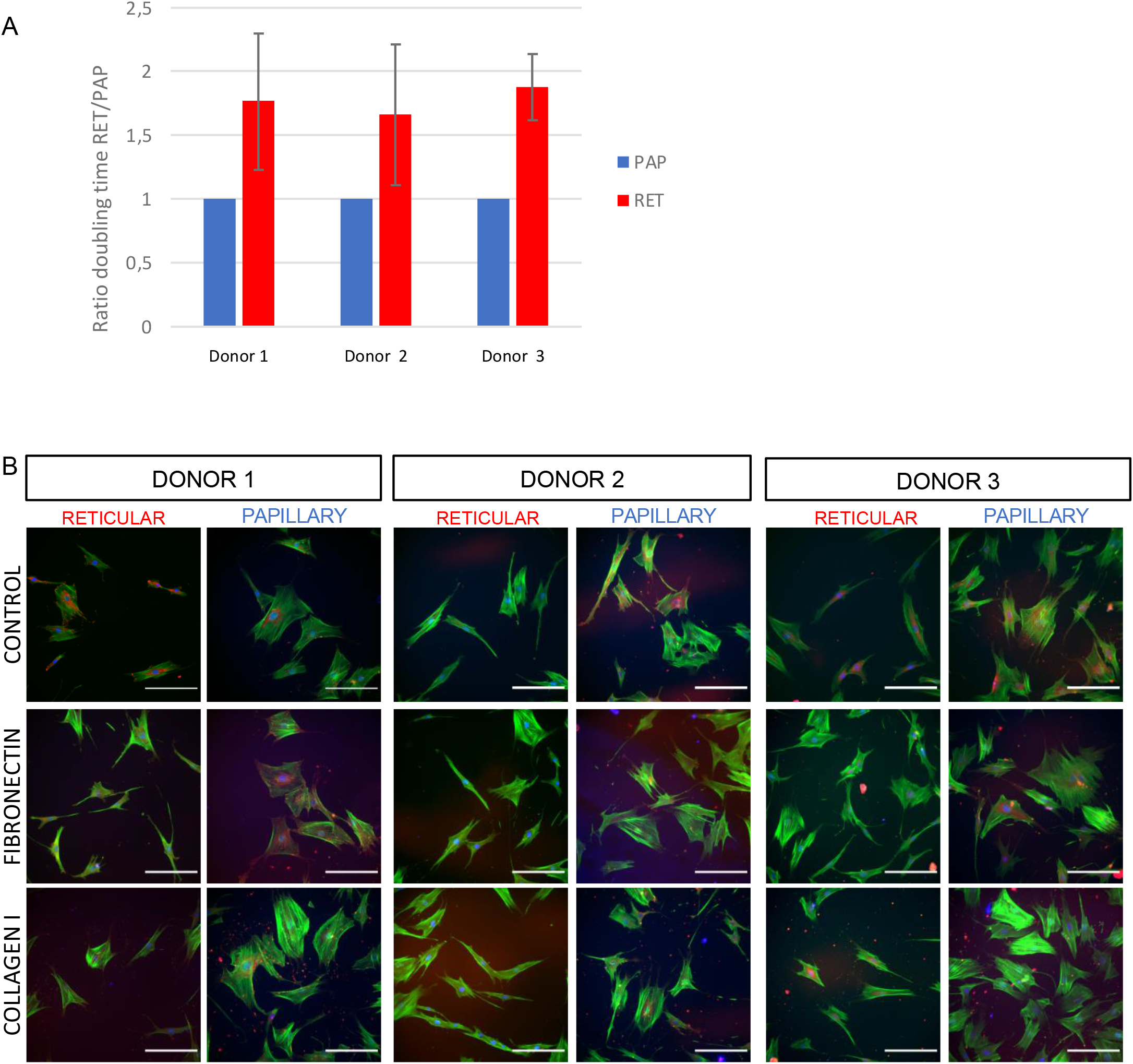
Validation of papillary and reticular features in the three donors used in this study. **A.** Difference in proliferation between fibroblasts populations was verified by calculating the ratio of doubling time between papillary and reticular fibroblasts. Site-matched fibroblasts were cultivated until one reached confluency and the ratio calculated for each donor. Graph represents the mean ratio of doubling time for at least six experiments ± SD. **B.** Morphology of isolated papillary and reticular fibroblasts observed by immunofluorescence for adhesion protein vinculin and cytoskeleton labelling with fluorescent phalloidin. Scale bar: 200 μm.

**Figure S2 :**
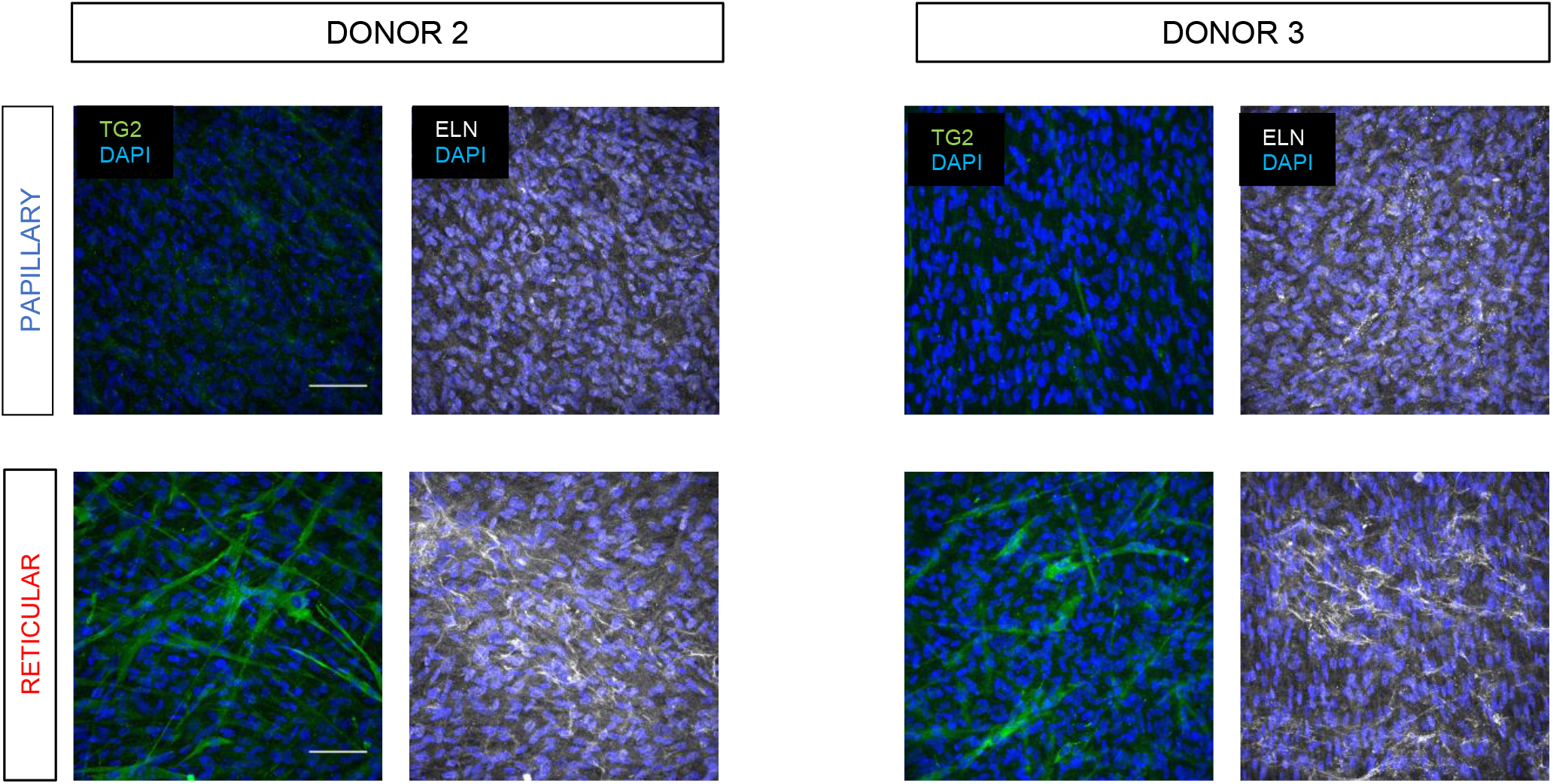
Expression of papillary and reticular matrisome markers in papillary and reticular cell sheets in the two other donors used in this study. Immunofluorescence stainin of elastin and TG2 expression in papillary and reticular cell sheets of donor 2 and 3 after two weeks of culture. Scale bar: 200 μm.

**Figure S3:**
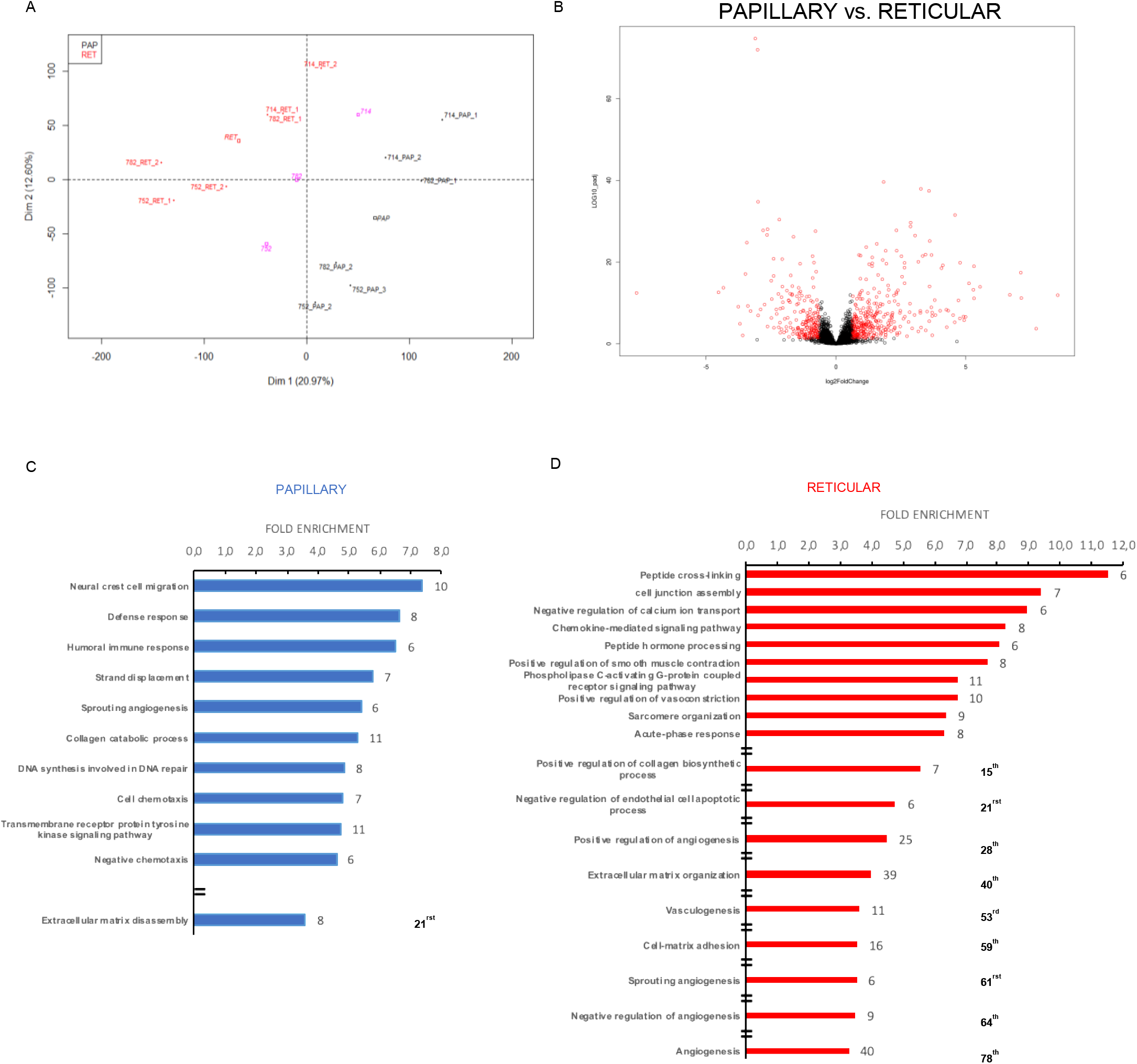
Bioinformatic analysis of papillary and reticular cell sheets transcriptomes. **A.** Principal component analysis (PCA) of full transcriptomes of papillary and reticular fibroblasts cultured as cell sheets. Dimension 1 is the one that segregates the more all samples (responsible for 20.97% of the variance) and cluster them in two groups that correspond to fibroblasts sub-populations. Samples are also separated according to the donor nature. **B.** Volcanoplot of all sequenced genes coding for proteins. Genes that are differentially regulated between papillary and reticular fibroblasts are indicated in red. FC>1.5 ; padj<0.05.Main enriched biological processes GO terms enriched in papillary (**C**) and reticular cell sheets (**D**). Go terms are displayed in decreasing order of fold enrichment for each population of fibroblasts. Gene counts are indicated for each GO term on the right hand side of the histogram. The tenth first GO terms are reported followed by all the GO terms related to blood vessel and ECM with their corresponding rank reported in bold. Biological Process Direct, Fold Enrichment ≥3, padj<1, gene count ≥5.

**Figure S4 :**
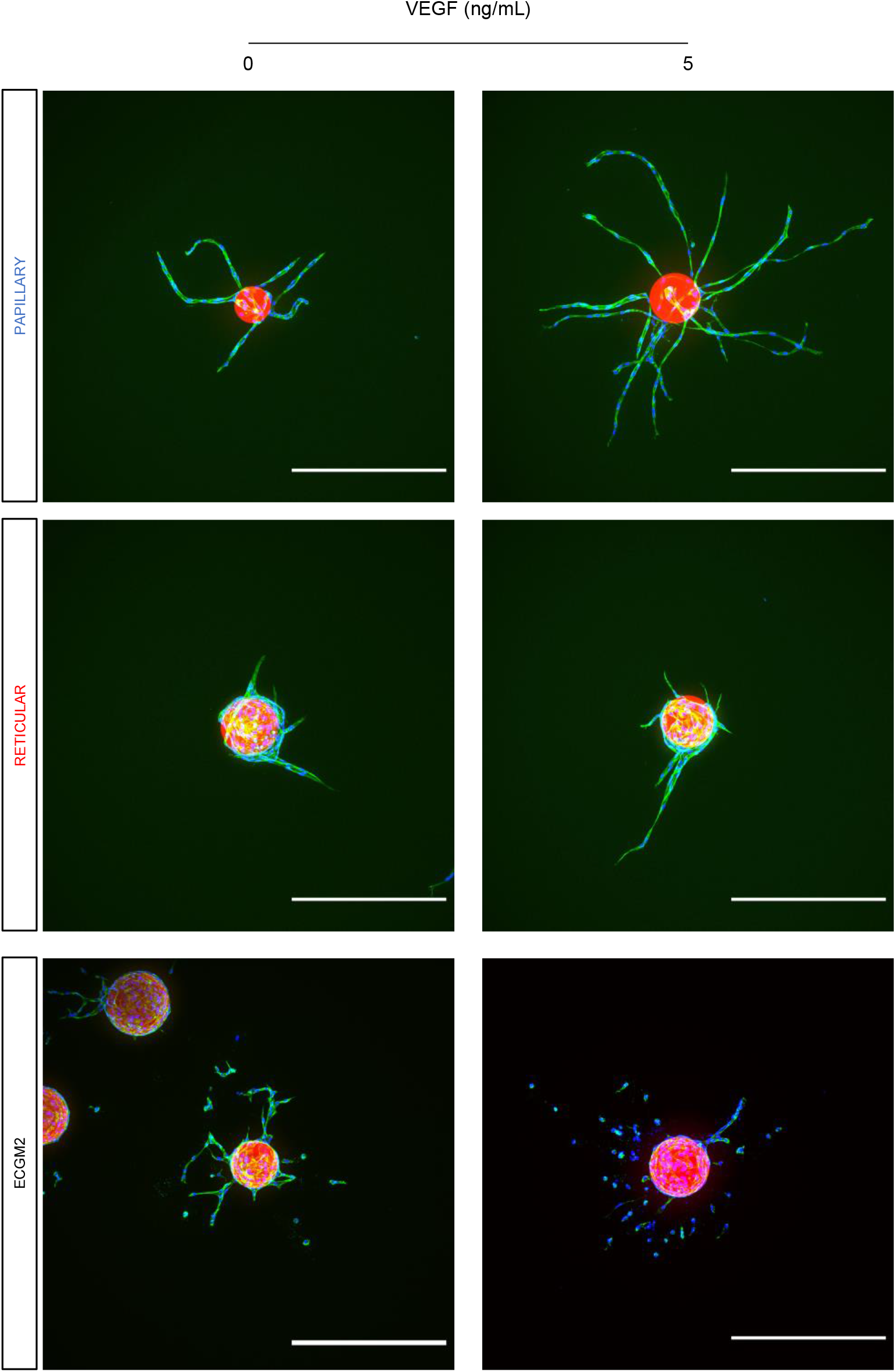
3D angiogenesis assay requires fibroblasts conditioned media independently from VEGF stimulation. Representative images of capillary formation in cytodex bead assay in the presence or absence of VEGF stimulation using capillary or reticular fibroblasts conditioned medium. Reticular fibroblasts allowed the formation of capillaries in the presence of VEGF to a lesser extent compared to papillary fibroblasts. The addition of endothelial medium ECGM2 did not allow capillary formation but induced the migration of endothelial cells from the bead. Maximum intensity projections. Scale bar: 500 μm.

